# Molecular markers of mechanosensation in glycinergic neurons in the avian lumbosacral spinal cord

**DOI:** 10.1101/2022.01.28.478253

**Authors:** Kathryn E. Stanchak, Kimberly E. Miller, Eric W. Lumsden, Devany Shikiar, Calvin Davis, Bingni W. Brunton, David J. Perkel

## Abstract

Birds are exceptionally adept at controlling their body position. For example, they can coordinate rapid movements of their body while stabilizing their head. Intriguingly, this ability may rely in part on a mechanosensory organ in the avian lower spinal cord called the lumbosacral organ (LSO). However, molecular mechanotransduction mechanisms have not been identified in the avian spinal cord. Here, we report the presence of glycinergic neurons in the LSO that exhibit immunoreactivity for myosin7a and epsin, molecules essential for function and maintenance of hair cells in the inner ear. Specifically, we find glycinergic cell bodies near the central canal and processes that extend laterally to the accessory lobes and spinal ligaments. These LSO neurons are reminiscent of glycinergic neurons in a recently-described lateral spinal proprioceptive organ in zebrafish that detects spinal bending. The avian LSO, however, is located inside a series of fused vertebrae called the synsacrum, which constrains spinal bending. We suggest the LSO may be a modification and elaboration of a pre-existing mechanosensory spinal network in vertebrates. A mechanistic understanding of its function may be an important clue to understanding the evolution and development of avian locomotion.

## 1 Introduction

Animals must be able to sense the position of their body to execute targeted movements, especially in response to environmental perturbations. Birds are often noted for their exceptional body-positioning abilities, including keeping their head still while their body twists and turns in response to wind or a swaying perch. As in other vertebrates, the vestibular organs of the inner ear and distributed proprioceptors affiliated with the musculoskeletal system provide sensory inputs to the avian body positioning control system. Interestingly, birds are also hypothesized to have a mechanosensory organ within their highly-modified lumbosacral spinal cord, known as the avian lumbosacral organ (LSO), which may provide additional mechanosensory information [11]. The LSO is found in all extant birds and may vary in morphology depending on a species’ locomotor habits [16]. However, the mechanotransduction mechanisms of the LSO remain unresolved.

It is becoming increasingly clear that vertebrates have spinal neurons that can provide sensory information on body positioning by detecting spinal bending. For instance, edge cells detect stretch at the lateral margin of the lamprey spinal cord [4]. Cerebrospinal fluid-contacting neurons (CSF-cNs) within the ependymal cells surrounding the central canal—found across vertebrates—sense alterations to CSF flow in the central canal due to spinal bending [1, 12]. Recently, a lateral glycinergic proprioceptive organ has been described in zebrafish that detects stretching of neural tissue near intervertebral discs as the spine bends [13].

While these examples of spinal mechanosensory neurons all detect bending of the spine, the avian LSO lies within the synsacrum, a rigid series of fused vertebrae that prevent the spinal cord from bending (Fig. 1A, B, C, D). Indeed, the LSO has a striking set of specializations that make it anatomically distinct from the rest of the spinal cord. First, a glycogen body sits within a dorsal split of the spinal cord. The central canal passes through the base of the glycogen body such that the ependymal cells surrounding the central canal are fully encapsulated by glycogen body tissue (Fig. 1D, E). Second, the spinal cord has segmentally-repeating, laterally-projecting accessory lobes, which are protrusions of neural tissue that roughly align with transverse, canal-like recesses in the surrounding bone (Fig. 1D). Modified spinal membranes route cerebrospinal fluid through these recesses [11]. Third, the entire neural structure is suspended within the synsacrum by a modified dentate ligament. At the ventrolateral corners of the spinal cord, the accessory lobes and the lateral dentate ligament adjoin the white matter (Fig. 1D, Fig. 2A, B, C). Based on these anatomical specializations, the LSO is hypothesized to work by transducing cerebrospinal fluid flow through mechanosensory neurons in the accessory lobes [10] or by acting as a preloaded vibratory structure [7].

**Figure 1:**
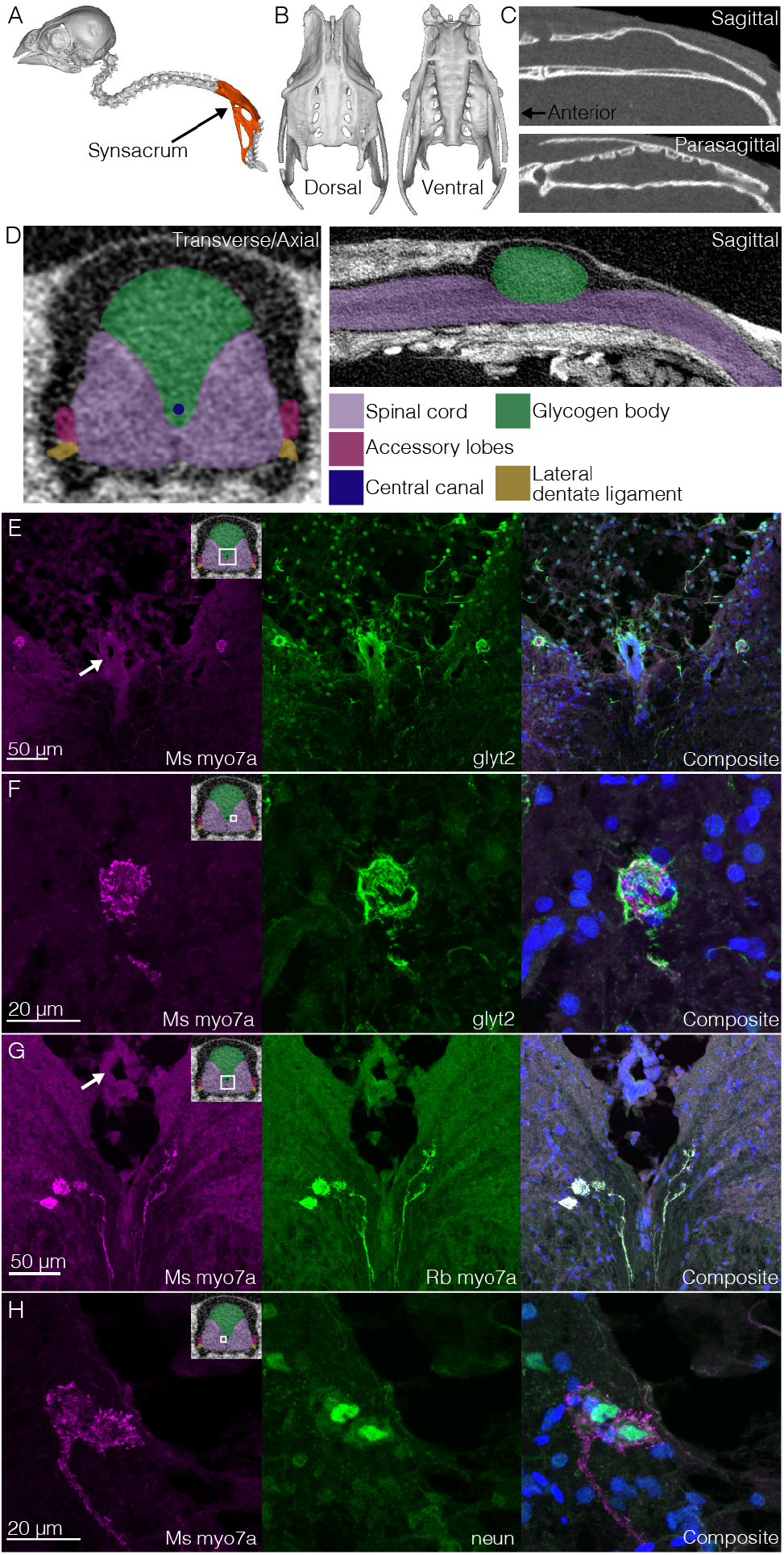
Glycinergic neuronal cell bodies near the the central canal of the avian lumbosacral organ (LSO) display myosin7a immunoreactivity. (A) The synsacrum (fused to the pelvic bones) is highlighted within a reconstructed CT scan of the zebra finch axial skeleton (Yale Peabody Museum specimen #125076). (B) Dorsal and ventral views of the synsacrum, reconstructed from the CT scan. (C) A sagittal section through the vertebral canal of the synsacrum CT scan shows the dorsal expansion of the vertebral canal that holds the glycogen body. A parasagittal section shows transverse canal-like structures on the internal dorsal side of the vertebral canal that roughly align with the accessory lobes (see [16]). (D) A diagram of the neural tissue of the LSO based on dissection and histology observations and a contrast-enhanced CT scan (data from [16]). (E) Cells near the central canal (middle) that are immunoreactive for both glyt2 and myo7a. (F) Higher-resolution image of the right cell from (E). (G) Labeling with two different myo7a antibodies (mouse and rabbit). (H) neun/fox3 labeling within the myo7a+ cells. White arrow in the first panel of subfigure (E) and (G) point to the ependymal layer surrounding the central canal. Upper right diagram in the first panel of subfigures (E-H) indicates approximately where in the transverse section the image was taken.

**Figure 2:**
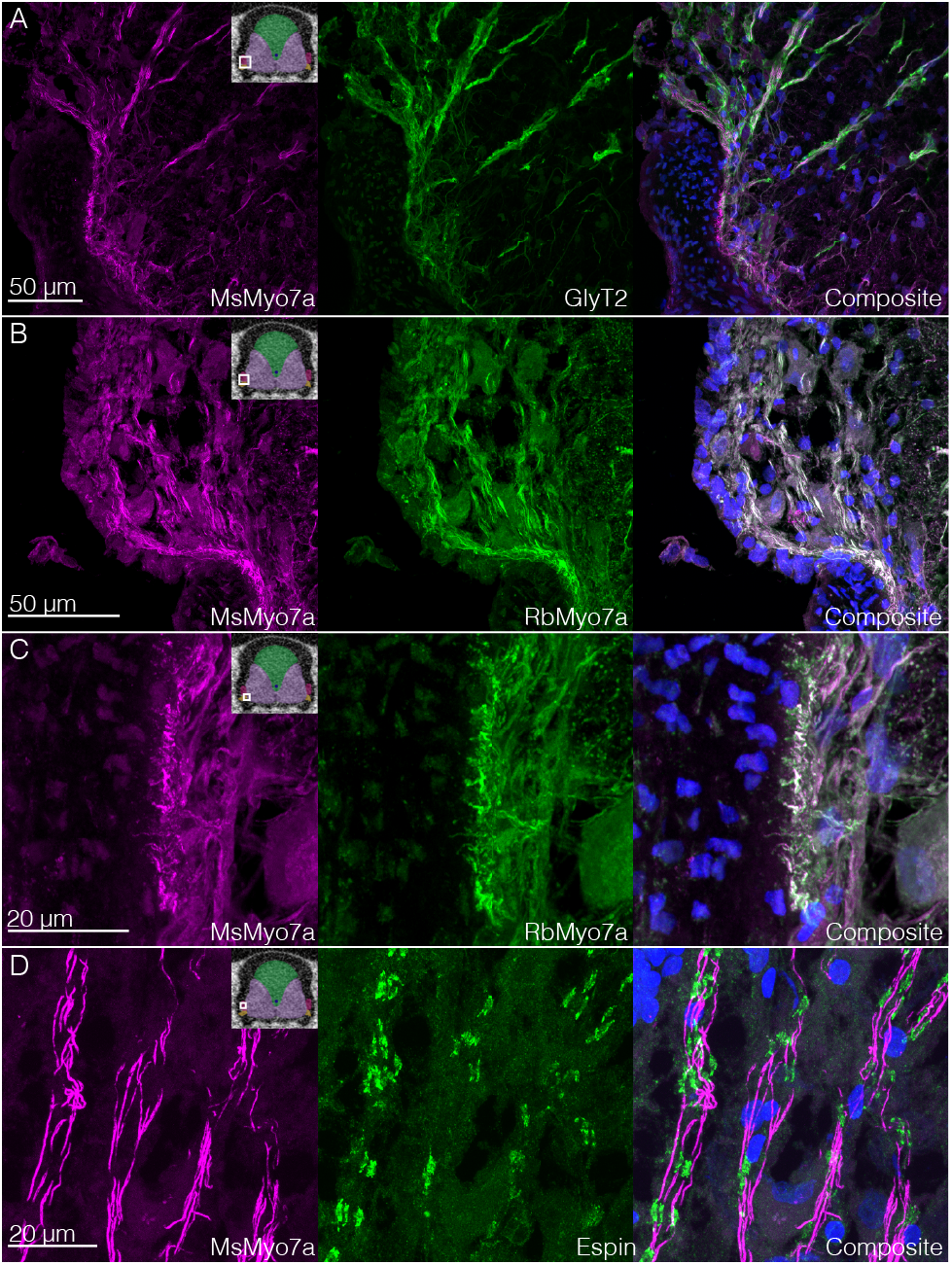
Lateral glycinergic processes are immunoreactive for proteins also present in inner ear hair cell stereocilia. (A) Glyt2+ ventrolateral processes are immunoreactive for myo7a. Margin of spinal cord with dentate ligament is at lower left; upper left is accessory lobe. (B & C) The accessory lobe processes (B) and terminations at the dentate ligament (C) double-label with two different antibodies for myo7a (mouse and rabbit). (D) Myo7a-immunoreactive processes in the accessory lobe also react with anti-espin. (AD) In composite images, blue is DAPI. Upper right diagram in the first panel of each subfigure indicates approximately where in the transverse section the image was taken.

While some cellular-level neuroanatomy of the LSO has been described, little is known about the molecular features of putative mechanosensory cells. In 1905, Imhof [5] illustrated cells and processes particular to the avian LSO, including around the central canal. Neurons within the accessory lobes produce a number of neurotransmitters, including glycine, glutamate, and GABA, as well as the enzyme choline acetyltransferase [9]. To date, no molecular mechanotransduction mechanism has been identified in the LSO.

Here, we describe neurons within the avian LSO that have molecular characteristics of known mechanosensory cells and are well positioned to transduce mechanical signals. These results are critical evidence supporting the hypothesis that the LSO has a role in proprioception and balance in birds.

## 2 Results

Immunohistochemical assays of the zebra finch LSO demonstrated medial neuronal cell bodies and ventrolateral processes that expressed the sodium- and chloride-dependent glycine transporter 2 (glyt2). The glycinergic medial cell bodies tended to be found near the central canal but not within the ependymal cell layer or the glycogen body that surrounds the central canal (Fig 1D and 1E). Further immunohistochemical assays demonstrated that these medial glycinergic cells were also immunoreactive for myosin 7a (myo7a, Fig. 1E, 1F, and 1G), an atypical myosin essential for proper inner ear hair cell function [8] and neun/fox3, a neuronal cell body marker (Fig 1H).

The glycinergic (glyt2+) ventrolateral processes traversed the gray matter to the white matter. Some processes extended into the accessory lobes, and others terminated at the margin between the dentate ligament and the spinal cord (Fig. 2A). These processes were also immunoreactive for myo7a (Fig. 2B and 2C). The processes within the accessory lobes were immunoreactive for both myo7a and espin antibodies, which in transverse sections colocalized in an alternating pattern (Fig. 2D). Espin is an actin-bundling protein, especially in sensory cells; [?, e.g.,]]sekerkova2011roles. Two different antibodies (from different manufacturers) for myo7a colabeled cellular structures (Fig. 1F, Fig. 2B and C). It was unclear from our sections whether these lateral processes extend from the medial cell bodies.

## Discussion

The putative mechanosensory neurons in the avian LSO described here comprise a lateral group of neurons that express glyt2, as in the lateral spinal proprioceptive organ of zebrafish [13]. We suggest these glyt2+ neurons in the avian LSO may be a modified version of the zebrafish lateral glycinergic proprioceptive organ. Notably, there is evidence these lateral mechanosensory zebrafish cells initially develop from progenitors near the central canal [13], which may be a developmental connection between the medial avian cells described here and the zebrafish cells.

There is no clear connection between CSF-cNs and the glyt2+ cells reported here. While CSF-cNs can be found ectopically (outside of the ependymal layer surrounding the central canal; [17, 6]), they are known to be GABAergic across vertebrates [3]. Further, a recent transcriptome analysis of spinal dynophin-lineage cells found a cluster with characteristics of CSF-cNs but comparatively low expression of glyt2 transcript (SLC6A5; Fig. 2 in [15]).

Our observations that glycinergic neurons in the LSO have immunoreactivity for myo7a and espin are critical evidence of their mechanosensory potential. Both myo7a and espin are found in inner ear hair cell stereocilia, and myo7a is directly implicated in mechanotransduction in hair cells [8, 14]. In addition, espin is also found in the cilia of CSF-cNs (as is the atypical myosin 3b; [2]).

The fused vertebrae of the bird synsacrum do not allow the lower spine to bend; therefore, the CSF-cNs and lateral proprioceptive spinal organ cannot function as they have been described in other vertebrates. Rather, the neuronal processes in the accessory lobes may sense cerebrospinal fluid flow (as hypothesized and summarized in [11]) while the processes at the junction between the spinal cord and the dentate ligament may detect differential vibration among tissue types (according to the vibration/suspension hypothesis in [7]).

Our finding of the presence of myo7a- and espin-immunoreactivity in LSO neurons is a starting point that critically enables testing of cellular, organ, and organismal-level mechanotransduction mechanistic hypotheses. Myo7a in particular may be a promising target for molecular manipulations. The ability of birds to impeccably control their body position may have helped them diversify into a clade with multifunctional locomotor specializations, including the capacity for terrestrial bipedalism as well as flight. Thus, continued research on the avian LSO may reveal foundational physiological and developmental mechanisms that enabled the evolution and diversification of avian locomotion.

## 3 Materials and Methods

### Tissue preparation

The spinal column was harvested from *(Taeniopygia guttata)* individuals used for other experiments. Most were post-fixed in 4% paraformaldehyde in phosphate buffer (PB) for 24h. Results from 7 individuals are reported here. Two spinal columns were alternatively harvested from transcardially-perfused individuals. In most samples, the LSO region of the spinal cord was dissected from the surrounding synsacrum bone and placed in 20% sucrose in PB for a minimum of overnight. The region of the LSO surrounding the glycogen body was then sectioned transversely at a cryostat in 4 alternating series with negative controls. For two samples (labeled with mouse-antimyo7a + rabbit-antiglyt2 and mouse-antimyo7a + rabbit-antimyo7a), the synsacrum bone was first decalcified using 12% EDTA neutralized with NaOH before the entire spinal column was sectioned. We saw no difference in reported results between decalcified and undecalcified specimens, but we note that anti-myo7a labeling seemed sensitive to extended time pre-freezing in other samples tested but not reported here.

### Immunohistochemistry

Sections were briefly dried then washed 3x in 0.1% Triton X-100 in phosphate buffered saline (PBST) before they were blocked with 5% neutral goat serum (Abcam or Vector Labs). Sections were then incubated for double-labeling experiments with primary antibody (AB) diluted in PBS, washed 3x in PBST, incubated with secondary AB, then washed in PBST, PBS, and dH20 and incubated for 10 minutes in 1:1000 aqueous DAPI (Biotium) prior to mounting with Fluoromount G (SouthernBiotech). Two timing protocols were used: a short protocol with a 1h block, overnight incubation with primary Ab, and 2h incubation with secondary Ab; and a long protocol with overnight block, 2 night incubation with primary Ab, and overnight incubation with secondary Ab. For some Abs we altered dilutions across experiments (noted below) but we saw no difference in reported results.

Primary Abs used were rabbit-antiglyt2 (Alomone AGT-012; 1:400 n=3 or 1:200 n=1; 13/14 immunogen amino acids identical to *T. guttata* glyt2 according to BLAST-P search), mouse-antimyo7a (DSHB 138-1, 1:100 n=3 or 1:200 n=4; avian cross-reactivity listed by manufacturer), rabbit-antimyo7a (Proteus 25-6790, 1:400 n=3 1:500 n=2; avian cross-reactivity listed by manufacturer), rabbit-antiespin (Invitrogen PA5-55941, 1:200 n=5; labeled interior side of ependymal cells in *T. guttata* spinal cord, where espin+ apical tips of CSF-cNs are located; 54/64 immunogen amino acids identical to *T. guttata* espin according to BLAST-P search), rabbit-antineun/fox3 (Invitrogen 702022; 1:200 n=3 or 1:500 n=2; labeled cells in spinal cord as expected). The double-labeling experiments were rabbit-antiglyt2 and mouse-antimyo7a (n=4); rabbit-antimyo7a and mouse-antimyo7a (n=5); rabbit-antiespin and mouse-antimyo7a (n=5); and rabbit-antineun and mouse-antimyo7a (n=5). Secondary Abs used were goat anti-mouse or goat anti-rabbit AlexaFluor 488, 568, and 647 (Invitrogen; 1:300). Slides were viewed and imaged on Nikon Eclipse epifluorescence and Olympus FV1000 confocal microscopes.

## Acknowledgements & Funding

We thank Claire Wyart for discussions about CSF-cNs and espin, as well as Elora Reilly for help with specimens. Yale Peabody Museum provided access to YPM orn 125076 CT scan data, the collection of which was funded by oVert TCN. The files were downloaded from www.MorphoSource.org, Duke University. Funding was provided by the Washington Research Foundation, the H. Stewart Parker Endowed Faculty Fellowship, and the Air Force Office of Scientific Research Award FA9550-19-1-0386 to BWB; and the University of Washington Department of Biology and Virginia Bloedel Hearing Research Center to DJP.

